# Gestational Chlorpyrifos Exposure Imparts Lasting Alterations to the Rat Somatosensory Cortex

**DOI:** 10.1101/2025.02.19.639085

**Authors:** Jeffrey A. Koenig, Catherine Haga, Nathan Cramer, Asaf Keller

## Abstract

Chlorpyrifos is an organophosphorus pesticide used extensively in agricultural and residential settings for nearly 60 years. Gestational, sub-acute exposure to chlorpyrifos is linked to increased prevalence of neurodevelopmental disorders. Animal studies have modeled these neurobehavioral detriments, however, the functional alterations in the brain induced by this exposure remain largely unknown. To address this, we used a rat model of gestational chlorpyrifos exposure to interrogate the alterations in the developing somatosensory (barrel) cortex. Rat dams were exposed to chlorpyrifos (5 mg/kg) or vehicle on gestational days 18-21 via subcutaneous injection, with no overt acute toxicity. Acetylcholinesterase was modestly inhibited but returned to baseline levels by postnatal day 12. We performed whole-cell patch clamp recordings on postnatal days 12-20 in both male and female progeny of the treated dams. A spike timing dependent plasticity protocol revealed changes to the normal development of use-dependent plasticity, including interference in long-term synaptic depression. Recording inhibitory synaptic activity revealed an increase in the frequency of spontaneous postsynaptic currents and in paired pulse ratios, in conjunction with a significant decrease in miniature postsynaptic currents. These findings suggest a presynaptic mechanism of inhibited GABA release, with potential disinhibition of inhibitory neurons. Evaluation of barrel cortex development displayed disruptions to normal barrel field patterning, with increases in both the septal area and total barrel field. We provide evidence for functional and structural alterations during brain development induced by *in utero* exposure to the organophosphorus pesticide chlorpyrifos that may account for the well-established behavioral outcomes.

**SIGNIFICANCE STATEMENT:** We demonstrate persistent alterations to synaptic function and plasticity in the somatosensory cortex following a brief, sub-acute exposure to the organophosphorus pesticide chlorpyrifos in a gestational rat model. These occur in conjunction with structural changes to cellular patterning of this brain region. These previously unknown consequences are potential causal mechanisms to the well-established neurodevelopmental detriments associated with early life exposure to chlorpyrifos. Clarifying these mechanisms could aid in ameliorating or preventing their persistent effects.

## INTRODUCTION

First characterized and produced in Germany during the 1930s-1940s, organophosphorus (OP) compounds have been ubiquitously employed as agricultural and residential insecticides ever since (Costa, 2018). Their primary mechanism of action is through potent and irreversible inhibition of acetylcholinesterase (AChE), the enzyme responsible for hydrolysis of acetylcholine at cholinergic synapses and neuromuscular junctions (Taylor, 2011). AChE is an evolutionarily conserved enzyme that can lead to unwanted toxicity in humans and other mammals when dysregulated. The toxidrome and standard treatment following acute exposure is well established and includes anticholinergics and oxime administration to reactive inhibited AChE (Marrs et al., 2006; Wiener and Hoffman, 2004). Of greater public health concern and increased scientific focus are the effects of environmental OP exposures that occur unbeknownst to the population. These exposures happen in the absence of overt toxicity or significant inhibition of AChE, potentially involving non-cholinergic mechanisms (Burke et al., 2017; Jaga and Dharmani, 2003). This concern is of special importance for exposures that occur during gestational development, as several large epidemiological studies have correlated this exposure with an increased prevalence of neurodevelopmental disorders, such as autism spectrum disorder and ADHD, and alterations to brain structure of children (Rauh et al., 2012; Schmidt et al., 2017; Shelton et al., 2014).

One of the most widely used and studied of these OP insecticides is chlorpyrifos (CPF). With strong evidence of neurodevelopmental toxicity, many countries have restricted its use, including a recent ban in the United States for food crops (EPA, 2021). Despite this, worldwide annual production and use of chlorpyrifos has increased from a reported 10,000 metric tons in 2007 to 50,000 metric tons in 2021 (ECHA, 2023). This continued and increasing utilization necessitates additional research investigating the potential mechanisms underlying the increased prevalence of neurodevelopmental disorders.

Several neurobehavioral rodent studies modeling sub-acute perinatal CPF exposure have demonstrated detriments to working memory and spatial learning, in addition to increased anxiety related behaviors, often in the absence of significant AChE inhibition (Aldridge et al., 2005; Levin et al., 2001; Mamczarz et al., 2016). *In vivo* investigations into neurochemical alterations outside of direct AChE inhibition have been limited. However, perinatal CPF exposure has been shown to significantly alter serotonergic signaling, with increased activity of receptor subtypes 5-HT_1A_ and 5-HT_2_ in a sex-dependent manner (Aldridge et al., 2004). Alterations to serotonergic signaling pathways could be of great importance as these play a critical role in cortical development and use-dependent plasticity (Lesch and Waider, 2012; Meunier et al., 2017; Miceli et al., 2013). Additionally, CPF can modulate endocannabinoid signaling pathways *in vivo* through inhibition of the endogenous endocannabinoid-hydrolyzing enzymes (Buntyn et al., 2017; Carr et al., 2011). Alterations to endocannabinoid signaling during development could have significant effects on normal cell proliferation, differentiation, and neuronal connectivity (Berghuis et al., 2007; Galve-Roperh et al., 2013).

The neurophysiological mechanisms underlying these neurobehavioral detriments remain unknown. We focus on the rat somatosensory (barrel) cortex as a model system to investigate the lasting effects of early CPF exposure. This well characterized cortical area processes information from the facial whiskers (Staiger and Petersen, 2021; Stüttgen and Schwarz, 2018). Sub-acute CPF exposure has been implicated in alterations in somatosensory network structure and function in both rodent models (Muller et al., 2014; Roy et al., 2004) and in humans (Silver et al., 2018; Van Wendel De Joode et al., 2016). Using an established late-gestational exposure paradigm (Garcia et al., 2002; Haviland et al., 2010; Levin et al., 2002), we studied changes to synaptic strength and plasticity in this region.

## MATERIALS AND METHODS

### Animals and treatments

All procedures adhered to the Guide for the Care and Use of Laboratory Animals and approved by the Institutional Animal Care and Use Committee at the University of Maryland School of Medicine. Male and female Sprague-Dawley rats were bred in our vivarium. Animals were housed under a standard 12/12 hour light/dark cycle and had *ad libitum* access to food and water. Pregnancy was confirmed by the presence of sperm in a vaginal lavage collection and set as gestational day (GD) 0. CPF (Chem Service, West Chester, PA) was dissolved in a 50/50 mixture of DMSO/peanut oil and injected subcutaneously (0.5 mL/kg) once daily on GD 18-21 at a dose of 5 mg/kg. Control dams received vehicle (DMSO/peanut oil) injections on the same schedule. Litters were culled to a maximum size of 10 by PND 3. Dams were pair-housed until GD 18. Both male and female pups (< PND 21) were used in all experiments at the ages listed in the results.

### Acetylcholinesterase Activity

Acetylcholinesterase (AChE) activity was measured using a modified Ellman colorimetric microplate assay described previously (Ellman et al., 1961; Shih et al., 2009). Forebrain, hindbrain, and heart tissue were collected on the day of birth (PND 0) and PND 2. The primary somatosensory cortex, one hemisphere of the cerebellum, and the heart apex were harvested on PND 12. AChE enzymatic activity was normalized to total protein using the bicinchoninic acid assay (Thermo Fisher Scientific, Waltham, MA).

### Slice Electrophysiology

*In vitro* slice electrophysiology was performed as we previously described (Alipio et al., 2021). Animals were deeply anesthetized with ketamine/xylazine, the brains were removed, and 300 μm coronal slices containing the primary somatosensory cortex were prepared. Slices were placed in a recording chamber continuously perfused (1.5 mL/min) with carbogen saturated artificial cerebrospinal fluid (ACSF) containing: 124 mM sodium chloride, 2.5 mM potassium chloride, 1.25 mM monosodium phosphate, 24 mM sodium bicarbonate, 12.5 mM glucose, 2 mM magnesium sulfate heptahydrate, and 2 mM calcium chloride dihydrate (Sigma-Aldrich, St. Louis, MO). Whole-cell patch clamp recordings of layer 2/3 pyramidal neurons were obtained in voltage clamp (-70 mV) mode. For EPSC recordings, the recording pipette (4-6 MΩ) was filled with a potassium gluconate solution containing: 120 mM potassium gluconate, 10 mM potassium chloride, 10 mM HEPES, 1 mM magnesium chloride, 0.5 mM EGTA, 2.5 mM magnesium ATP, and 0.2 mM GTP-Tris (Sigma-Aldrich, St. Louis, MO). For IPSC recordings, 6-cyano-7-nitroquinoxaline-2,3-dione (CNQX; 20 mM; Sigma-Aldrich, St. Louis, MO) and DL-2-amino-5-phosphonopentanoic acid (APV; 50 mM; Sigma-Aldrich, St. Louis, MO) were included in the ACSF and a high-chloride pipette solution was used containing: 70 mM potassium gluconate, 60 mM potassium chloride, 10 mM HEPES, 1 mM magnesium chloride, 0.5 mM EGTA, 2.5 mM magnesium ATP, 0.2 mM GTP-Tris. (Sigma-Aldrich, St. Louis, MO). Tetrodotoxin (1 μm; Bio-Techne, Minneapolis, MN) was additionally added to the ACSF for mIPSC recordings. Series resistance was monitored in all recordings by measuring the current evoked by a -5 mV square pulse at 30 second intervals.

The spike timing dependent plasticity (STDP) protocol was modified from Itami and Kimura (2012). Briefly, layer 2/3 pyramidal neurons were recorded in bridge mode with potassium gluconate pipette solution described above. Single action potentials were evoked by current injection (<1 nA; 10 ms) through the recording pipette. Electrical stimulation (<0.2 mA; 100 μs) was delivered through a bipolar electrode placed in layer 4, directly below the recorded cell, to produce a single-component EPSP with an amplitude of 3-10 mV. The pairing protocol consisted of 90 post-before-pre pairings, with an action potential delivered 25 ms prior to the EPSP every 7.5 seconds. EPSP amplitudes were recorded every 7.5 seconds for 5 minutes prior (baseline) and 40 minutes following the pairing protocol. Input resistance was monitored throughout the recording by measuring the change in potential induced by a -10 mA current injection. The post-pairing amplitude was calculated from the average amplitude between minutes 40 and 50.

### Histology

The barrel field was stained using cytochrome oxidase histochemistry using a modified protocol as previously described (Iwasato et al., 1997). Animals were deeply anesthetized via intraperitoneal injection of ketamine/xylazine and transcardially perfused with ice-cold PBS followed by 10% neutral buffered formalin (NBF). The brains were extracted, and the cortex was separated. The cortex was flattened between glass slides with spacers (∼2 mm), and post-fixed overnight at room temperature with 10% NBF. 90 μm thick horizontal sections were prepared with a vibratome. Staining solution was prepared with 1.6 g sucrose, 20 mg diaminobenzidine, and 40 mg cytochrome C (Sigma-Aldrich, St. Louis, MO) in 40 mL of PBS. Sections were incubated with staining solution at 37°C in an incubator shaker for 4-5 hours, followed by an additional 18 hours at room temperature. Following a PBS rinse, sections were mounted on charged slides and allowed to dry overnight. Sections were dehydrated, cleared, and coverslipped with DPX mounting media (Sigma-Aldrich, St. Louis, MO). Barrel maps were analyzed using a modified method from Heidrich et al (2020). Briefly, barrels from rows C-E and columns 1-3 of the posteromedial barrel subfield were traced by contrast threshold in ImageJ (NIH). The centroids of the border barrels were calculated and connected with lines. The septal area was contained within this connected perimeter. The border of the map area consisted of the border of the barrels and septa area as depicted in Figure 8.

For analysis of neuron morphology, cells were filled with 0.1% biocytin (Thermo Fisher Scientific, Waltham, MA) during electrophysiology recordings. Slices were fixed in 10% NBF for 24-48 hrs following experiments and then stored in PBS until processing. Slices were incubated for 24 hrs in streptavidin conjugated Cy3 (1:1000; Jackson ImmunoResearch Laboratories, West Grove, PA) then mounted with aqueous media. Dendritic arbors were traced and analyzed using Imaris 10 filament tracer (Oxford Instruments, Concord, MA). A separate Sholl analysis was performed on the apical and basal dendritic regions by counting the intersections across concentric circles spaced 10 µm apart.

### Statistical Analysis

Statistical analyses were conducted with GraphPad Prism 10 (Boston, MA). Statistical significance was set as *p* < 0.05. Parametric tests were used if the appropriate assumptions were met. Otherwise, nonparametric tests were used. The statistical tests used are listed in each figure legend. There were no differences between rats of different litters in the reported endpoints thus they were combined for analysis. Sex differences were investigated where powered and noted in Results.

## RESULTS

### Effect of gestational CPF exposure on dam and litter health

We first determined if repeated daily administration of CPF (5 mg/kg) during GD 18-21 (Fig 1A) produces overt toxicity by measuring litter size and dam/pup weights. We weighed pregnant dams daily during the 4-day treatment period (Fig 1B). Vehicle and CPF treated dams gained weight at the same rate during the treatment period. Litter sizes were indistinguishable between CPF (median: 12.5 CI 11-14) and vehicle (median: 15 CI 12-15) treated groups (Fig 1C). Pup weight at time of birth (PND 0) showed no difference between treatment groups (Fig 1D), with a mean weight of 6.6g for both vehicle and CPF. Pup weights measured throughout the experimental period (Fig 1E) similarly demonstrated no difference between CPF (R^2^ = 0.90) and vehicle (R^2^ = 0.90) treated groups. These data demonstrate a lack of overt toxicity induced by our gestational CPF exposure model.

**Figure 1.**
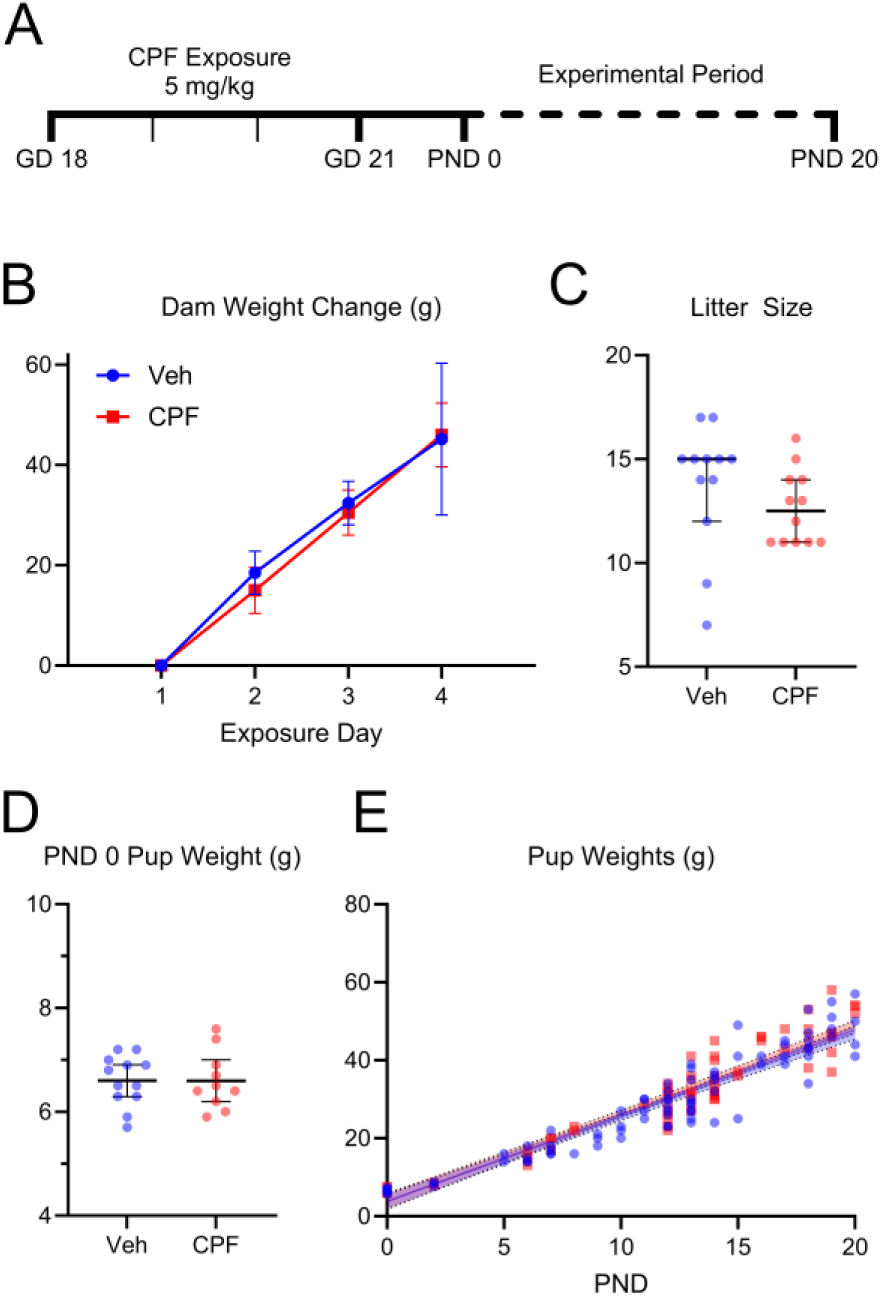
Gestational chlorpyrifos exposure does not affect dam or litter health. **A.** Timeline of gestational CPF exposure and experimental period. **B.** Daily weight change of pregnant rat dams (vehicle n=12 and CPF n = 12) during the experimental treatment period (GD 18-21) was similar between groups (F (1, 21) = 0.27, p = 0.61, mixed-effects analysis). **C.** Litter size (vehicle n = 12 and CPF n = 12) was unaffected by CPF exposure (U = 44, p = 0.10, Mann Whitney test). **D.** Pup weight at birth (PND 0; vehicle n = 12 and CPF n = 10) was similar in both treatment groups (t(18) = 0, p > 0.99, Welch’s unpaired t test). **E.** Pup weight gain pre-weaning (PND 0-20; vehicle n = 89 and CPF n = 82) was similar in both CPF and vehicle treated groups (F(2,267) = 0.98, p = 0.38, extra-sum-of-squares F test). Mean ± 95% CI (**B, E**), median ± 95% CI (**C**), best-fit line and 95% CI (**D**).

### Inhibition of AChE activity

Our exposure model further aimed to inhibit AChE activity only moderately and transiently, as inhibition of >50% AChE activity in the brain is associated with acute fetal impairments (Qiao et al., 2002). Measurements of AChE enzymatic activity at PND 0 (24hrs after final CPF exposure; Fig 2A) showed inhibition in the forebrain (73.1 ± 7.6%), hindbrain (69.6 ± 8.0%), and heart (42.1 ± 9.5%) compared to vehicle treated animals. AChE activity in the CPF group remained inhibited at PND 2 (Fig 2B), measured in the forebrain (76.6 ± 6.4%), hindbrain (79.7 ± 4.7%), and heart (60.4 ± 8.0%). At PND 12 (Fig 2C), AChE activity in CPF treated animals had recovered, as measured in the cortex (88.5 ± 13.1%), cerebellum (96.6 ± 10.1%), and heart (97.5 ± 5.0%).

**Figure 2.**
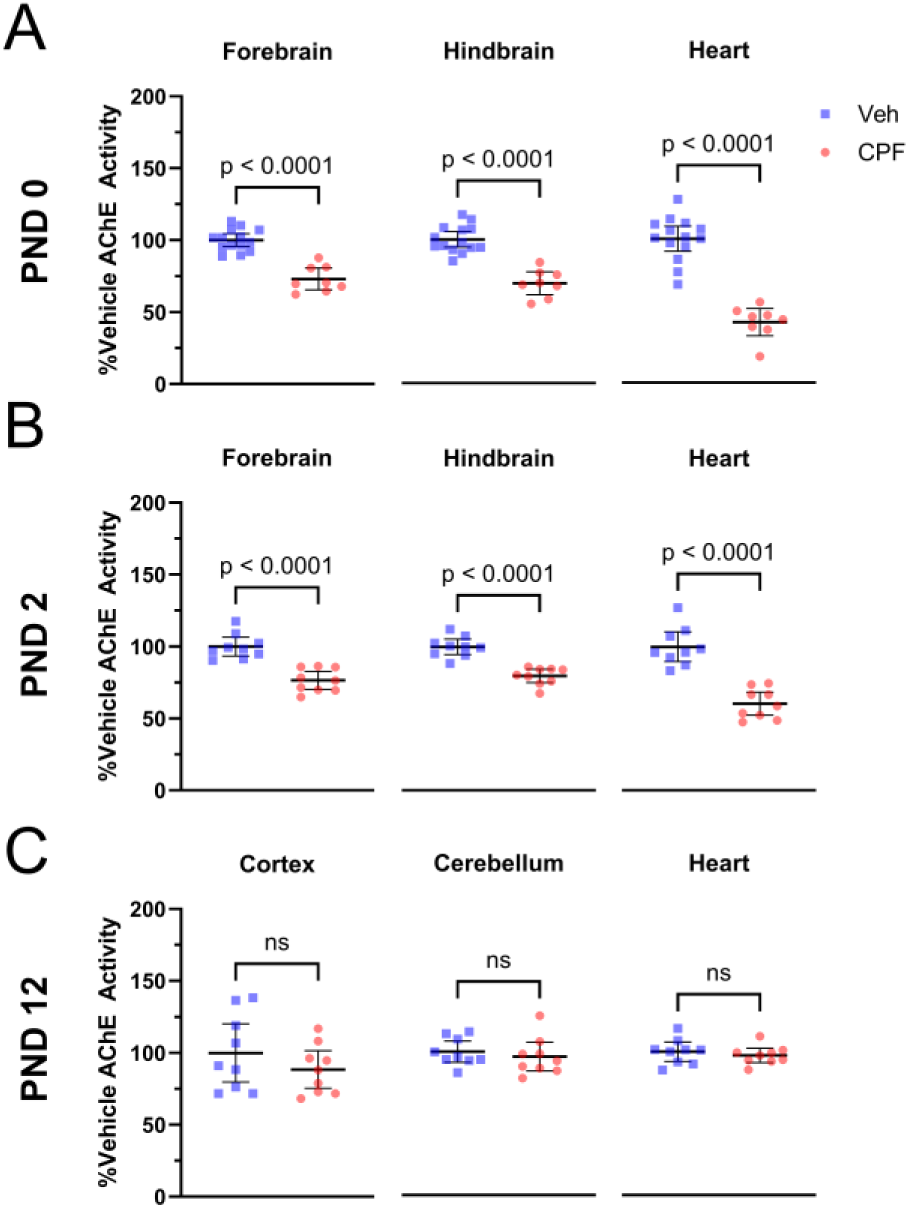
Gestational chlorpyrifos exposure produces moderate and transient inhibition of AChE activity in offspring. **A.** At PND 0, AChE activity (vehicle n = 14 animals from 6 litters and CPF n = 8 animals from 4 litters) of the CPF exposed group, compared to vehicle, was inhibited in the forebrain (t(12.6) = 7.1, p < 0.0001), hindbrain (t(14.6) = 7.3, p < 0.0001), and heart (t(18.3) = 10.1, p < 0.0001, Welch’s unpaired t test). **B.** At PND 2 (vehicle n = 9 animals from 3 litters and CPF n = 9 animals from 3 litters), AChE activity in the CPF exposed group remained inhibited in the forebrain (t(16.0) = 5.9, p < 0.0001), hindbrain (t(15.7) = 6.6, p < 0.0001), and heart (t(15.1) = 7.0, p < 0.0001, Welch’s unpaired t test). **C.** AChE activity at PND 12 (vehicle n = 9 animals from 3 litters and CPF n = 9 animals from 3 litters) was the same in both the CPF and vehicle exposed groups measured in the cortex (t(13.7) = 1.1, p = 0.29), cerebellum (t(14.8) = 0.63, p = 0.54), and heart (t(14.8) = 0.7, p = 0.49, Welch’s unpaired t test).

### Intrinsic neuronal properties

To determine if CPF treatment results in lasting changes in neuronal membrane properties, we obtained whole-cell patch recordings from layer 2/3 pyramidal neurons in the somatosensory (barrel) cortex (Fig 3A), from PND 12-20 aged animals. Intrinsic excitability was determined through current injection in 25pA steps from 0 to 200pA. There was no measured difference in the firing rate-current relationship between neurons from vehicle and CPF exposed animals (Fig 3B). Measurements of rheobase (62.5 ± 16.3 pA; 75.6 ± 15.9 pA), resting membrane potential (-67.2 ± 3.1 mV; -67.6 ± 2.2 mV), half-width (2.2 ± 0.1 ms; 2.1 ± 0.1 ms), spike height (41.7 ± 2.5 mV; 43 ± 2.3 mV), and threshold potential (-36.7 ± 3 mV; -36.1 ± 1.7 mV) also revealed no difference between vehicle and CPF exposed animals, respectively (Fig 3C).

**Figure 3.**
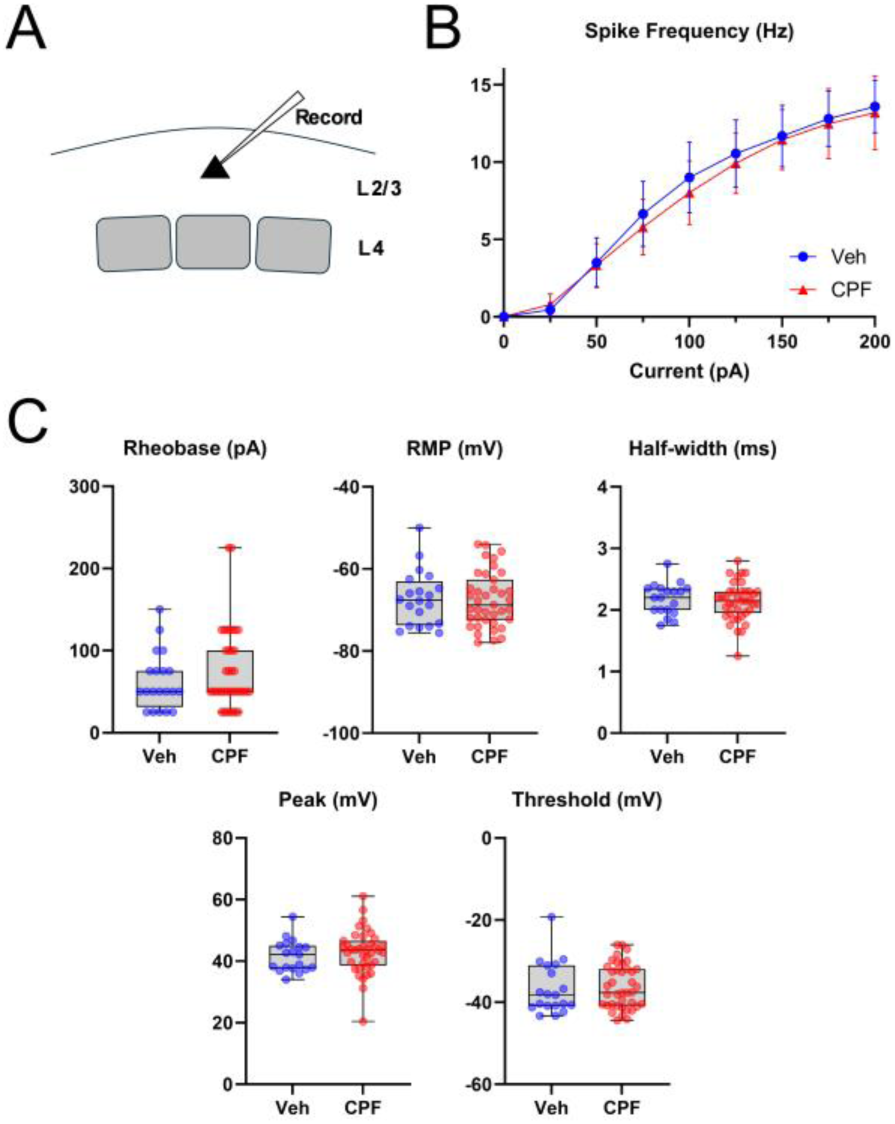
Gestational chlorpyrifos exposure has no effect on the intrinsic excitability of pyramidal neurons in the somatosensory (barrel) cortex. **A.** Schematic of patching location of layer 2/3 pyramidal neurons in the barrel cortex. **B.** Firing rate-current (F-I) recordings (vehicle n=20 cells from 12 animals and CPF n=39 cells from 20 animals) reveal no difference between groups (F (1, 57) = 0.20, p = 0.65, mixed-effects analysis). **C.** Measurement of rheobase (U = 341.5, p = 0.43, Mann Whitney test), resting membrane potential (RMP; t(38.5) = 0.22, p = 0.83), half-width (t(42.4) = 0.58, p = 0.57), peak height (t(47.9) = 0.78, p = 0.44), and threshold potential (t(31.4) = 0.36, p = 0.72, Welch’s unpaired t test) similarly demonstrated no difference between vehicle and CPF treated animals (number of cells/animals as in A). Mean ± 95% CI (**B**), box plots with whiskers of minimum and maximum values (**C**).

### Excitatory synaptic activity

There is increasing evidence for the role of dysregulation in the excitatory/inhibitory synaptic balance in neurodevelopmental disorders such as ADHD and autism spectrum disorder (Ferranti et al., 2024; Gao and Penzes, 2015). To investigate whether gestational CPF is imparting such an imbalance, we examined if CPF treatment results in lasting changes in glutamatergic synaptic transmission by recording spontaneous excitatory postsynaptic currents (sEPSCs) from layer 2/3 pyramidal neurons in the barrel cortex. Example traces from CPF and vehicle exposed animals are shown in figure 4A. The mean frequency of sEPSC events (Fig 4B) was similar between vehicle (3.34 ± 1.01 Hz) and CPF (3.13 ± 0.63 Hz) exposed groups. Similarly, there was no difference in mean sEPSC amplitudes (Fig 4C) between the vehicle (12.97 ± 0.47 pA) and CPF (12.61 ± 0.28 pA) exposed groups. No sex differences were observed in these experimental outcomes, so data from males and females were combined. These findings indicate that gestational CPF exposure has no lasting effect on basal glutamatergic transmission in the barrel cortex.

**Figure 4.**
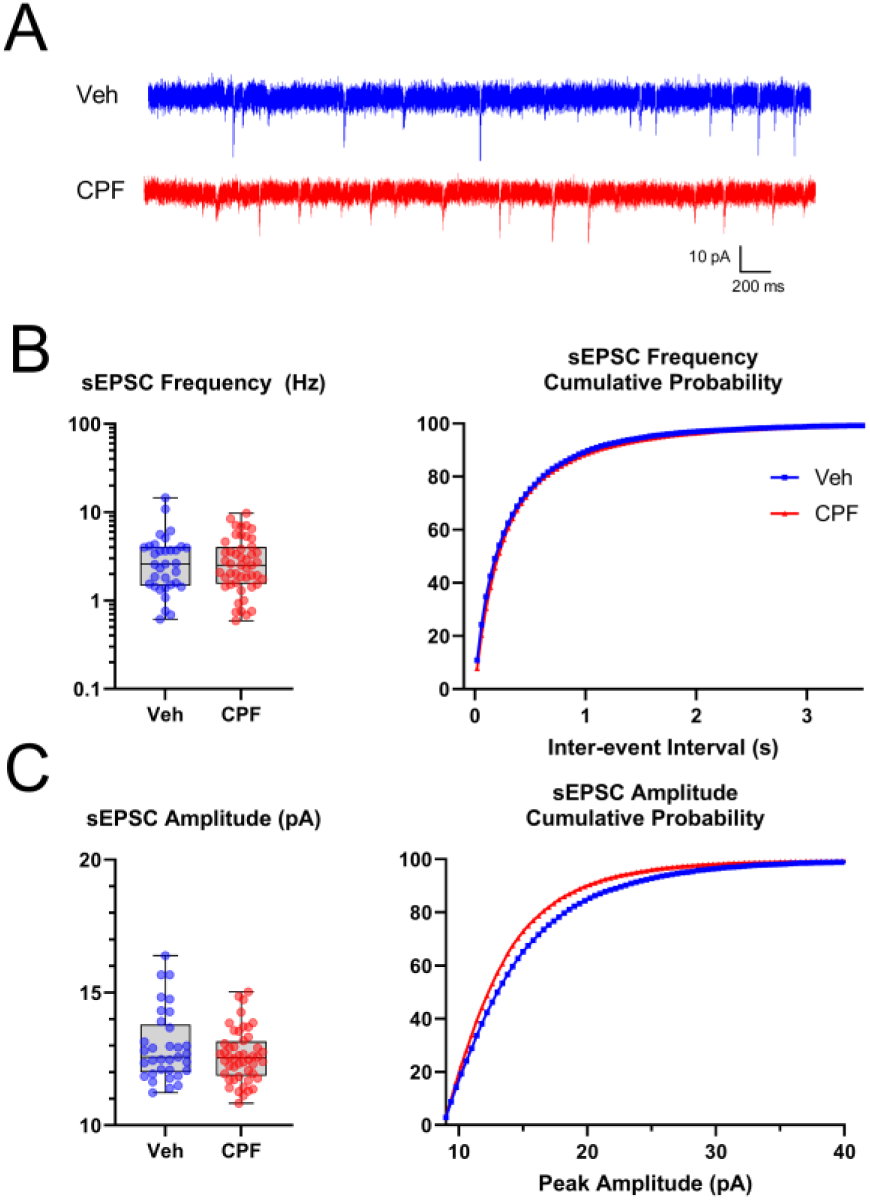
Gestational chlorpyrifos exposure does not impact excitatory synaptic activity in the barrel cortex. **A.** Example traces of sEPSCs from vehicle and CPF exposed animals. **B.** Neither the frequency (t(56.6) = 0.36, p = 0.72, Welch’s unpaired t test; D = 0.04, p > 0.99, Kolmogorov-Smirnov test) nor amplitude (**C**; t(55.5) = 1.3, p = 0.19, Welch’s unpaired t test; D = 0.13, p = 0.21, Kolmogorov-Smirnov test) of sEPSC events were different between vehicle and CPF exposed animals (vehicle n = 33 cells from 23 animals and CPF n = 48 cells from 25 animals). Box plots with whiskers of minimum and maximum values (**B, C**), cumulative probability histogram (**B, C** curves).

### Inhibitory synaptic activity

Having seen no effect on excitatory transmission, we next assessed inhibitory GABAergic synaptic activity through measurement of spontaneous inhibitory postsynaptic currents (sIPSCs). Figure 5A depicts example recordings from CPF and vehicle exposed animals. The mean sIPSC frequency (Fig 5B) was approximately 84% higher in neurons from the CPF group (2.73 ± 0.88 Hz), compared to those from vehicle-treated animals (1.48 ± 0.41 Hz). There was a corresponding leftward shift in the interevent interval cumulative probability of the CPF group, compared with vehicle. There were no differences in mean sIPSC amplitudes (Fig 5C) between vehicle (21.31 ± 3.63 pA) and CPF (22.94 ± 3.05 pA) treated groups. No sex differences were observed in these experimental outcomes, so data from males and females were combined.

**Figure 5.**
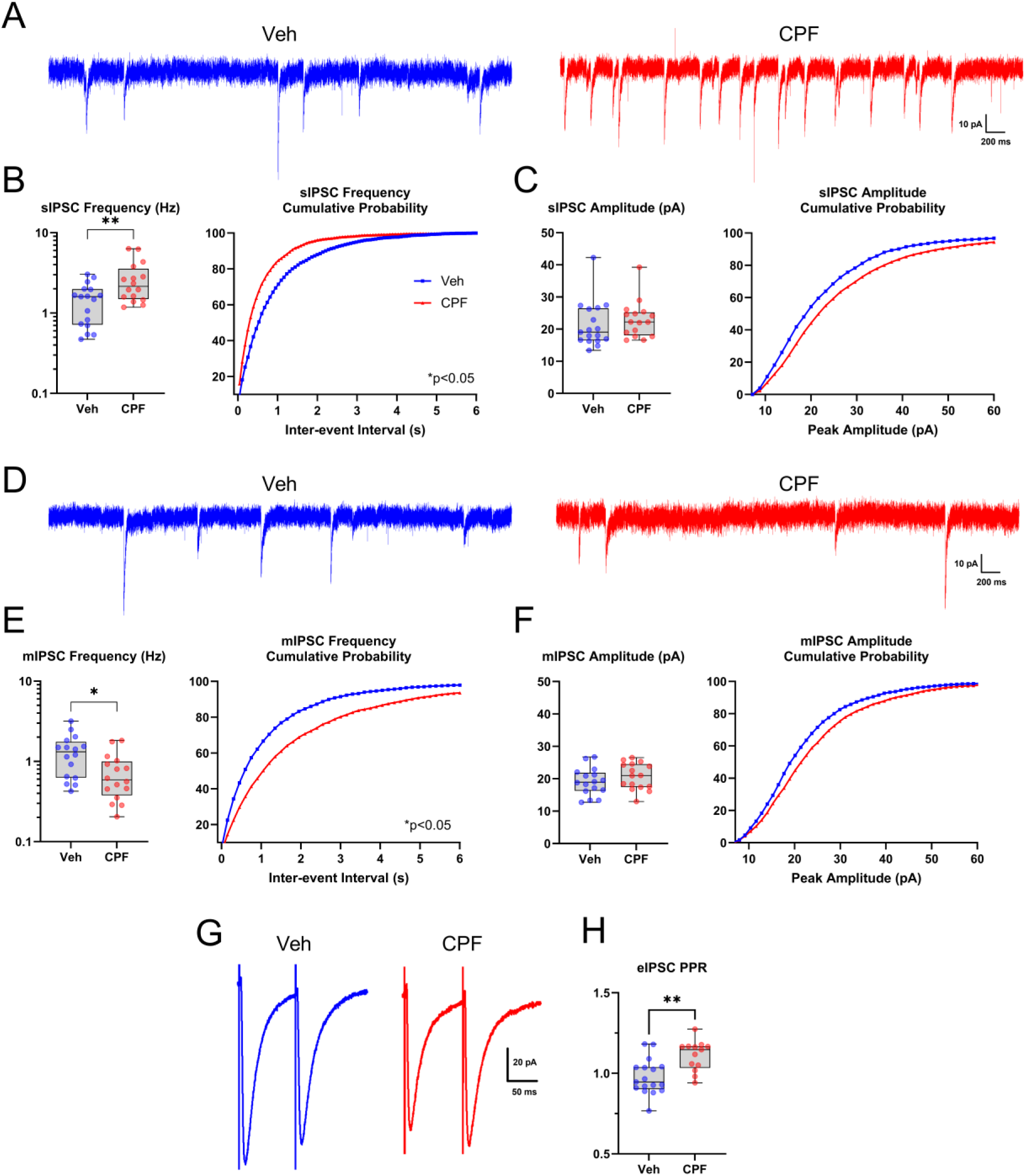
Gestational chlorpyrifos exposure alters GABAergic synaptic activity in the barrel cortex. **A.** Example traces of sIPSCs from vehicle and CPF exposed animals. **B.** The frequency of sIPSC events (vehicle n=17 cells from 8 animals and CPF n=16 cells from 8 animals) was higher in CPF exposed animals when compared to vehicle (t(21.4) = 2.7, p = 0.01, Welch’s unpaired t test), with a leftward shift in the cumulative probability curve (D = 0.24, p = 0.01, Kolmogorov-Smirnov test). **C.** The amplitude of sIPSC events showed no corresponding difference (t(30.4) = 0.73, p = 0.47, Welch’s unpaired t test; D = 0.15, p = 0.29, Kolmogorov-Smirnov test). **D.** Example traces of mIPSCs from vehicle and CPF exposed animals. **E.** The frequency of mIPSC events (vehicle n=16 cells from 7 animals and CPF n=16 cells from 10 animals) was lower in the CPF exposed group compared to vehicle (t(25.4) = 2.6, p = 0.02, Welch’s unpaired t test), with a rightward shift in the cumulative probability curve (D = 0.24, p = 0.009, Kolmogorov-Smirnov test). **F.** There was no difference in the amplitude of mIPSC events (t(30) = 1.17, p = 0.25, Welch’s unpaired t test; D = 0.18, p = 0.10, Kolmogorov-Smirnov test). **G.** Example traces of paired-pulse evoked IPSCs (eIPSCs). **H.** The PPR (vehicle n = 17 cells from 9 animals and CPF n = 13 cells from 5 animals) was greater in CPF exposed animals when compared to vehicle (t(27.6) = 3.6, p = 0.001, Welch’s unpaired t test). Box plots with whiskers of minimum and maximum values (**B, C, E, F, H**), cumulative probability histogram (**B, C, E, F** curves).

We also compared TTX-insensitive miniature inhibitory postsynaptic currents (mIPSCs) in CPF and vehicle animals, with example traces seen in figure 5D. The mean mIPSC frequency (Fig 5E) in the CPF group (0.75 ± 0.26 Hz) was approximately 44% lower compared to vehicle-treated (1.34 ± 0.41 Hz), with a rightward shift in the interevent interval cumulative probability. The mean amplitude of mIPSC events (Fig 5F) was similar in CPF (20.78 ± 2.18 pA) and vehicle (19.06 ± 2.24 pA) groups. These data suggest a decreased probability of presynaptic GABA release in the CPF exposed animals.

To further clarify this mechanism, we compared evoked inhibitory postsynaptic current (eIPSC) responses to paired pulse stimulation and calculated a paired pulse ratio (PPR), the ratio of the amplitude of the second pulse divided by the amplitude of the first pulse (Kim and Alger, 2001). A PPR above one is associated with a low probability of vesicle release (i.e., “weaker” synapses), whereas a PPR below one is associated with a high probability of release. Example paired-pulse responses, evoked by stimulating directly below the recorded neuron in layer 4, from CPF and vehicle exposed animals are shown in figure 5G. The mean PPR (Fig 5H) was 14% higher in the neurons from CPF (1.11 ± 0.06) exposed animals, compared to neurons from vehicle (0.98 ± 0.06) treated animals. These findings further suggest that CPF exposure results in lasting suppression of presynaptic GABA release.

### Spike timing dependent plasticity

Spike timing dependent plasticity (STDP) is a form of associative synaptic plasticity in which the temporal order of the presynaptic and postsynaptic action potentials determines the sign of plasticity – whether long-term synaptic depression (LTD) or potentiation (LTP) is induced (Feldman, 2012; Markram, 2011). This form of Hebbian plasticity is believed to underlie learning and information storage, as well as the development and refinement of neuronal circuits during brain development (Bi and Poo, 2001; Dan and Poo, 2006; Sjöström et al., 2008). Between the second and third week of rodent development, STDP of the L4 – L2/3 synapse in S1 undergoes use-dependent metaplasticity. Before this age, all temporal pairings of pre and postsynaptic action potentials result in LTP, whereas at older ages both LTP and LTD can be evoked, depending on the order and timing of pre and postsynaptic stimulation (Itami et al., 2016; Itami and Kimura, 2012). Here, we asked whether CPF treatment results in lasting changes in this metaplasticity.

We used a negative timing post-before-pre (-25ms) pairing protocol while recording, in bridge mode, from barrel cortex pyramidal neurons of PND 12-20 animals (Fig 6A). That is, a postsynaptic spike was evoked, by intracellular current injection, 25ms before evoking a postsynaptic EPSP by extracellular stimulation (see Methods). Figures 6B and 6C show example EPSP waveforms before and after the pairing protocol and the time course of changes in EPSP amplitudes demonstrating LTD and LTP induction, respectively. The stability of membrane resistance (Rm) throughout the recordings is shown below each graph.

**Figure 6.**
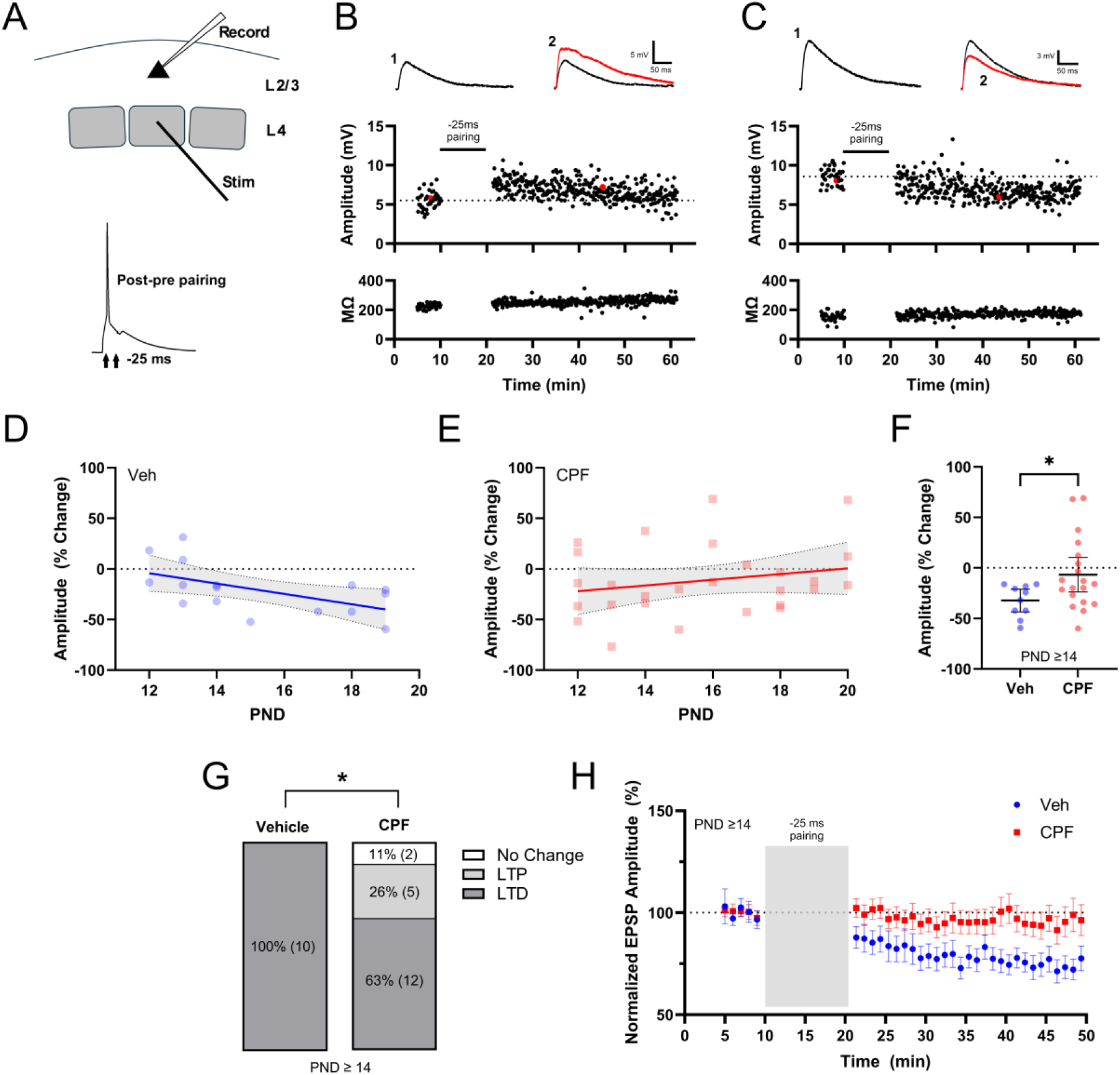
Gestational chlorpyrifos exposure alters use-dependent plasticity in the barrel cortex. **A.** Schematic of recording location of layer 2/3 pyramidal neuron in the barrel cortex with electrical stimulation of layer 4. Trace demonstrates example post-before-pre stimulation. **B.** Example of LTP and LTD (**C**) induction following post-before-pre pairing protocol with representative EPSP waveforms at baseline (1) and following pairing (2). The location of the representative traces in the data set are highlighted in red. **D.** The developmental progression of increasing LTD (vehicle n = 16 cells from 14 animals and CPF n = 27 cells from 25 animals) is altered in CPF exposed animals (**E**) when compared to vehicle (F(1,39) = 4.6, p = 0.04, extra-sum-of-squares F test). **F.** The % change in amplitude for animals ≥ PND 14 (vehicle n=10 cells from 9 animals and CPF n=19 cells from 17 animals) was diminished in CPF exposed animals compared to vehicle (t(26.6) = 2.7, p = 0.01, Welch’s unpaired t test). **G.** A greater proportion of cells (number of cells/animals as in F) were unable to demonstrate LTD in the CPF exposed group when compared to vehicle (p = 0.03, one-sided Fisher’s exact test). LTP was defined as an EPSP amplitude increase above 5% while LTD was defined as an amplitude decrease below -5%. **H.** The time-course of EPSP amplitude change following pairing was altered in the CPF exposed group compared to vehicle (F(1,27) = 4.4, p = 0.045, mixed-effects analysis). Best-fit line and 95% CI (**D, E**), mean ± 95% CI (**F, H**).

Consistent with previous reports, our pairing protocol evoked an increasing magnitude of LTD (Fig 6D) correlated with age (R^2^ = 0.31, p = 0.025) in the vehicle group. In contrast, gestational CPF exposure disrupted this progression (Fig 6E), such that a mix of LTP and LTD was evoked in different neurons across ages (R^2^ = 0.05, p = 0.263), altering the plasticity/age relationship compared to vehicle. Focusing specifically on ages after the previously established transition period (≥ PND 14), neurons from vehicle treated litters had a mean EPSP amplitude reduction of 32.3 ± 11.3%, which was diminished in CPF-treated animals, with a mean EPSP amplitude reduction of 6.6 ± 17.1% (Fig 6F). The fraction of neurons which produced LTD vs those that did not (Fig 6G), was also reduced after CPF exposure. The mean time-course of depression following pairing (Fig 6H) additionally demonstrated alterations, with lower LTD induction in neurons from CPF exposed animals. Taken together, these data demonstrate marked alterations in use-dependent metaplasticity in the barrel cortex induced by gestational CPF exposure.

### Cell morphology

Sub-acute OP exposure has been previously reported to alter neuron morphology measured as reduced dendritic branching in hippocampal pyramidal neurons (Mullen et al., 2016; Narasimhamurthy et al., 2023). To determine if gestational CPF exposure induces such changes in neuron morphology, we analyzed dendritic branching of barrel cortex layer 2/3 pyramidal neurons (Fig 7A). Sholl analysis revealed no difference in dendritic complexity between the vehicle and CPF exposed groups in either the apical (Fig 7B) or basal (Fig 7C) compartments. Additional measurements of total dendritic length and branch point count demonstrated no difference between groups in either the apical or basal compartments.

**Figure 7.**
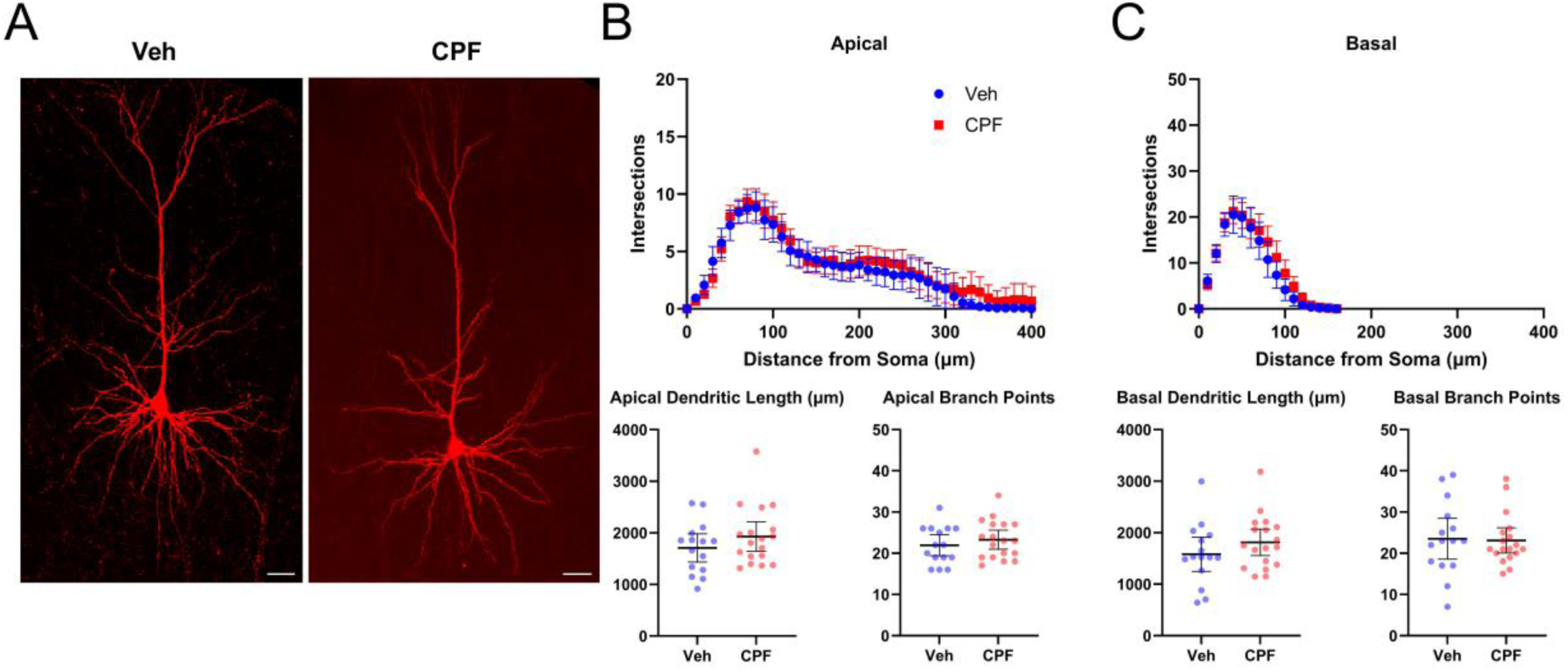
Gestational chlorpyrifos exposure does not affect pyramidal cell morphology in the barrel cortex. **A.** Example of biocytin-filled layer 2/3 pyramidal neurons from vehicle and CPF exposed animals. **B.** In the apical region, Sholl analysis revealed no differences between vehicle and CPF exposed animals (F(1,31) = 1.19, p = 0.28, two-way ANOVA). There was similarly no difference in total dendritic length and number of branch points (t(30.9) = 1.19, p = 0.24; t(30.1) = 0.83, p = 0.41, Welch’s unpaired t test). **C.** For basal dendrites, there was no difference in either Sholl analysis (F(1,31) = 1.4, p = 0.25, two-way ANOVA) or total dendritic length and number of branch points (t(27.9) = 1.18, p = 0.25; t(24.1) = 0.15, p = 0.88, Welch’s unpaired t test) between vehicle and CPF groups (vehicle n = 15 cells from 12 animals and CPF n = 18 cells from 15 animals). Mean ± 95% CI (**B, C**). Scale bar is 25 μm.

### Barrel Field

The rodent barrel cortex forms a somatotopic map where each facial whisker is directly related to a discrete structure (barrel) within layer 4 of the primary somatosensory cortex. Both molecular mechanisms and peripheral input underlie the proper organization of this barrel field early in development (Li and Crair, 2011; Petersen, 2007), with altered patterning being associated with lasting impairments to normal sensory processing (Miceli et al., 2017; Papaioannou et al., 2013; Su et al., 2021). Additionally, perinatal toxin exposure has been reported to detrimentally alter the organization of the barrel cortex (Chappell et al., 2007; Powrozek and Zhou, 2005; Wilson et al., 2000). We sought to determine if gestational CPF exposure interferes with this proper patterning. A tangential section of the barrel cortex displaying the complete posteromedial barrel subfield is shown (Fig 8A). Morphological analysis (Fig 8B) revealed a larger barrel map area (red outline) in CPF exposed animals (0.83 ± 0.10 mm^2^) when compared to vehicle (0.71 ± 0.06 mm^2^). Similarly, septa area (white outline) in the CPF group were larger (0.18 ± 0.04 mm^2^) compared to vehicle (0.11 ± 0.04 mm^2^). However, there was no difference in total barrel area between vehicle (0.58 ± 0.07 mm^2^) and CPF (0.64 ± 0.07 mm^2^) treated groups. These data demonstrate alterations to proper barrel field patterning, specifically a larger septa area, induced by gestational CPF exposure.

**Figure 8.**
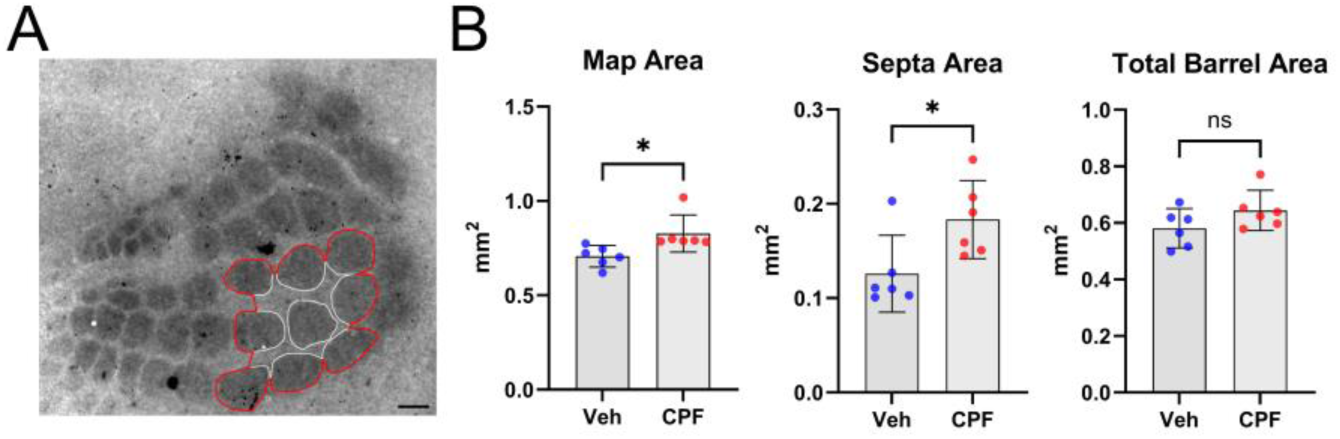
Gestational chlorpyrifos exposure disrupts barrel field patterns in the somatosensory cortex. **A.** Example cytochrome C oxidase-stained barrel field from vehicle treated PND 6 rat pup. The measured region of the barrel field (map area) is outlined in red, with individual barrels outlined in white. **B.** Map (t(8.1) = 2.72, p = 0.03) and septa (t(10) = 2.54, p = 0.03), but not total barrel area (t(10) = 1.62, p = 0.14, Welch’s unpaired t test), are increased in CPF exposed animals compared to vehicle (vehicle n = 6 and CPF n = 6). Mean ± 95% CI (**B**). Scale bar is 200 μm.

## DISCUSSION

Early life exposure to the organophosphorus pesticide, chlorpyrifos (CPF), is associated with an increased incidence of neurodevelopmental disorders in children (Bouchard et al., 2011; Rauh et al., 2012, 2006; Schmidt et al., 2017; Shelton et al., 2014). In rodent models, exposure to CPF results in lasting neurobehavioral deficits (reviewed in Todd et al., 2020). Here, we sought to identify functional alterations to cortical neurons, and the networks in which they are embedded, that may be driving these lasting neurobehavioral deficits. Using a brief, sub-acute exposure to CPF during gestation, we demonstrate that this exposure is associated with lasting alterations to inhibitory synaptic strength, disruptions to the developmental progression of use-dependent plasticity, and morphological changes to the somatosensory (barrel) cortex of the rat.

### AChE activity

Our model aimed to produce an asymptomatic gestational exposure with mild and transient inhibition of acetylcholinesterase (AChE) to mimic an agricultural or residential exposure of humans to CPF (Eaton et al., 2008; Reiss et al., 2015). As anticipated, there was no acute toxicity to the dam or litter, as there were no changes to litter size or pup/dam weight throughout the experimental period. There was a modest inhibition of AChE in the brain and peripheral tissue measured at PND 0, at levels below acute fetal toxicity (Qiao et al., 2002). Enzymatic activity quickly recovered to baseline levels by PND 12. The rapid recovery of AChE activity reported here agrees with previous reports in neonatal rodents (Pope et al., 1991; Song et al., 1997). This recovery of AChE activity prior to our electrophysiological recordings is significant, as increased cholinergic signaling due to persistently inhibited AChE would alter measurements of synaptic strength and the induction of spike timing dependent plasticity (Brzosko et al., 2019; Fuenzalida et al., 2021; Yang et al., 2014).

### Sex differences

Reports on neurobehavioral sex differences in models of CPF exposure are variable, likely due to variability in exposure periods. Females typically demonstrate higher sensitivity with earlier gestational exposures and males show more sensitivity with postnatal exposure paradigms (Aldridge et al., 2005; Haviland et al., 2010; Levin et al., 2002, 2001). Most previous reports associating prenatal CPF exposure with neurodevelopmental disorders in humans did not find sex-specific sensitivities (Eskenazi et al., 2007; Schmidt et al., 2017). Those that did reported a stronger association in males than females (Fortenberry et al., 2014; Marks et al., 2010). We did not to detect sex differences in the endpoints studied here. As our results looked exclusively at ages ≤ PND 20, we cannot exclude the possibility that sex differences become apparent at later ages (Imhof et al., 1993; Premachandran et al., 2020).

### Lasting effects on inhibitory synapses

Late gestational exposure to CPF led to an increase in the frequency of spontaneous inhibitory postsynaptic currents (sIPSCs) in layer 2/3 pyramidal neurons, with no concurrent change in the amplitude. Both findings are consistent with enhanced presynaptic GABA release (Zucker and Regehr, 2002). These inhibitory inputs most likely arise from parvalbumin (PV) interneurons, as they provide dense inputs and powerful inhibition to all neighboring cortical pyramidal cells (Packer and Yuste, 2011). Conversely, we discovered a decrease in the frequency of miniature inhibitory postsynaptic currents (mIPSCs) and an increase in the paired-pulse ratio (PPR) in CPF exposed animals. Both findings are consistent with suppressed GABA release by interneurons (Glasgow et al., 2019). These seemingly contradictory data could potentially be explained by increased spontaneous firing of the PV neurons.

Most inhibitory inputs to cortical PV neurons arise from other PV neurons and somatostatin expressing interneurons (Kubota et al., 2016; Xu et al., 2013; Yang et al., 2016). A reduction in presynaptic GABA release, as demonstrated in our mIPSC and PPR results, could result in disinhibition of PV neurons and an increase in their spontaneous activity (Laaris et al., 2000; Xiang et al., 1998). Enhanced firing by these fast-spiking interneurons could overcome a presynaptic inhibitory mechanism at the PV-pyramidal synapse and result in the increased sIPSC frequency reported here. A potential mechanism driving this presynaptic inhibition could be altered 5-HT signaling. Following gestational CPF exposure, binding to 5-HT receptors 5-HT_1A_ and 5-HT_2_ is permanently upregulated (Aldridge et al., 2004, 2003; Slotkin and Seidler, 2005). While the 5-HT_1A_ receptor has inhibitory presynaptic activity, this is primarily as an autoreceptor on 5-HT neurons (Altieri et al., 2013; Verge et al., 1985). The related receptor 5-HT_1B_ is highly expressed in the cortex, preferentially on presynaptic axon terminals, and its activation strongly suppresses both GABAergic and glutamatergic neurotransmitter release (Boschert et al., 1994; Bramley et al., 2005; Bruinvels et al., 1994; Hashimoto and Kita, 2008; Laurent et al., 2002; Puig and Gulledge, 2011). Future research may identify cell type-specific altered activity of the 5-HT_1B_ receptor following early CPF exposure.

Dysregulation of PV neuron activity is altered in several neurodevelopmental disorders, such as schizophrenia and autism spectrum disorder (Ferguson and Gao, 2018; Filice et al., 2020; Selten et al., 2018). PV neurons are key in regulating the activity of cortical networks (Freund, 2003; Nahar et al., 2021). In the barrel cortex, PV neurons have a unique role in shaping receptive field size and response dynamics (Kwegyir-Afful and Keller, 2004; Kyriazi et al., 1996; Miller, 2001). Proper control of receptive field size is critical for proper sensory perception (Kolasinski et al., 2017). PV neurons also regulate cortical gamma oscillations, the rhythmic fluctuations in local field potentials that control connectivity between brain regions and facilitate perception, cognition, and memory (Buzsáki and Wang, 2012; Fries, 2009; Sohal et al., 2009). Disruption to these oscillations have been associated with various neurodevelopmental and psychiatric disorders (Bitzenhofer, 2023; Guan et al., 2022; Traub, 2010; Yener and Başar, 2013). The role early CPF exposure has on potential PV neuron disfunction and the causative relationship to the established neurobehavioral detriments modeled in rodents remain to be investigated.

### Lasting effects on synaptic plasticity

Use-dependent plasticity is the process by which sensory experience or stimulation can regulate the formation and maturation of neural circuits, typically presented as long-term potentiation (LTP) or long-term depression (LTD) (Chaudhury et al., 2016; Ganguly and Poo, 2013; Katz and Shatz, 1996). The coordinated pairing of presynaptic stimulation with postsynaptic spiking can induce this plasticity in what is known as spike timing dependent plasticity (STDP) (Bi and Poo, 2001; Feldman, 2000). In STDP, the temporal order of presynaptic stimulation and postsynaptic action potential determines whether LTD or LTP is induced (Caporale and Dan, 2008). These forms of Hebbian plasticity underlies learning and information storage, as well as the development and refinement of neuronal circuits during brain development (Dan and Poo, 2006, 2004).

In the cortex, the propensity of circuits to undergo LTP or LTD undergoes a developmental switch. In the layer 4 to layer 2/3 synapse of the barrel cortex, the pathway activated by whisker stimulation (Armstrong-James et al., 1992), plasticity switches from LTP only to a bidirectional LTP/LTD response around the third postnatal week (∼PND 14) (Itami and Kimura, 2012). Here we performed a post-before-pre stimulation, with layer 2/3 spiking proceeding layer 4 stimulation. At the ages studied here (PND 12-20) this would typically induce increasing magnitudes of LTD, which was apparent in our vehicle treated animals (Fig 6). In contrast, in CPF exposed animals, there was a mix of LTP and LTD across the entire age range, completely ablating the normal progression to bi-directional plasticity during this critical period. This anomaly may have lasting effects on the proper development and maturation of this circuit (Brzosko et al., 2019; Desai et al., 2006). The proper activity of both PV neurons and 5-HT signaling are critical for the regulation of this plasticity (Cavaccini et al., 2018; Higa et al., 2022; Kimura and Itami, 2019; Vickers et al., 2018), lending further evidence for their potential disruption following early CPF exposure.

### Lasting changes in barrels structure

Evaluation of barrel field patterning revealed significant alterations, particularly increases in septa width and total barrel map area. Proper formation of the barrel field is sensitive to increased 5-HT activity (Cases et al., 1996), with activation of the 5-HT_1B_ receptor preventing the refinement of thalamocortical axons into discrete barrels (Rebsam et al., 2002). The altered barrel patterning described here represents a potential disruptive mechanism for the proper organization and refinement of the somatosensory cortex, which may underlie associated neurodevelopmental disorders (Cascio, 2010; Khan et al., 2015; Marco et al., 2011; Nair et al., 2013).

### Conclusions

Our study describes lasting functional and structural alterations to the somatosensory cortex induced by gestational CPF exposure. The disruptions to the excitatory/inhibitory balance, use-dependent plasticity and thalamocortical organization are all associated with the occurrence of the same neurodevelopmental disorders seen following early CPF exposure. Additional studies clarifying these effects could offer novel biomarkers or potential therapeutics following this early life exposure to CPF.

